# Second impact of an acute total body irradiation on thymic epithelial cells

**DOI:** 10.1101/2024.07.30.605780

**Authors:** Kano Namiki, Wataru Muramatsu, Takahisa Miyao, Hiroto Ishii, Rin Endo, Naho Hagiwara, Maki Miyauchi, Masaki Yoshida, Nobuko Akiyama, Taishin Akiyama

## Abstract

The thymus, a crucial organ for T cell development, undergoes transient involution following exposure to sublethal total body irradiation. The impact of sublethal irradiation on the thymus is reportedly bimodal: thymic involution recurs after the recovery of the thymus from the initial impact of acute sublethal irradiation. While the second impact of acute irradiation has been acknowledged for thymocytes, its influence on thymic epithelial cells (TECs), which are crucial for thymic T cell differentiation and selection, remains to be elucidated. In this study, we aim to elucidate this influence. Mice were subjected to acute sublethal total body irradiation, and TECs were evaluated at three distinct time points: during the initial impact, during the recovery phase post-initial impact, and during the second impact phase. Flow cytometry analysis revealed that during the second impact phase, mTECs were reduced, whereas cTECs remained unaffected. Among mTECs, the subset expressing high levels of the co-stimulatory molecule CD80 (mTEC^hi^) experienced the most pronounced reduction. RNA sequencing analysis of mTEC^hi^ cells at early differentiation stages revealed significant alterations in gene expression profiles during the second impact phase. Notably, gene signatures of tuft-like TECs and Aire-expressing TECs were preferentially influenced in these mTEC^hi^ subpopulations. These findings suggest that acute total body irradiation disrupts mTEC frequencies and gene expression in a bimodal manner, possibly compromising thymic functions over extended periods.

## Introduction

In the intricate process of T cell differentiation in the thymus, thymic epithelial cells (TECs) emerge as pivotal orchestrators^1^. TECs, classified into cortical TECs (cTECs) and medullary TECs (mTECs) based on their localization, assume distinct roles. Cortical TECs primarily facilitate the initial differentiation of thymocytes and the positive selection of self-MHC-restricted T cells. Conversely, medullary TECs exhibit a unique ability to express and present a diverse array of self-tissue-restricted antigens (TSAs)^2^, crucial for governing the negative selection of self-reactive T cells and fostering the development of thymic regulatory T cells. The expression of TSAs is uniquely regulated by the autoimmune regulator (AIRE)^3^ of which malfunction has been implicated in the onset of autoimmunity in both human and murine systems ^4^. This intricate interplay underscores the fundamental importance of TECs in shaping the immune landscape and warrants comprehensive investigation.

Recent studies using single-cell analysis have unveiled significant heterogeneity among TECs^5–10^. In addition to the simple categorization of cTECs and mTECs, the latter exhibit diverse subtypes including Aire-expressing mTECs (Aire^+^ mTECs), CCL21-expressing mTECs, transit-amplifying mTECs, and Post-Aire mTECs. The Post-Aire mTEC is the differentiation stage following Aire^+^ mTEC, consisting of mimetic TEC subpopulations that possess chromatin structures and gene expression profiles similar to various tissue cells^10^. Thus, these TEC subtypes and subpopulations exhibit unique gene expression profiles, which contribute to the diversity of TSAs and the distinct functions in the thymus^10,11^.

The thymus undergoes both aging-dependent and -independent involution processes^12–16^. Aging-dependent thymic involution is typically irreversible. Conversely, aging-independent involution can be triggered by stressors such as psychological stress, viral infections, chemicals, and irradiation, and it can be homeostatically recovered^14,16–18^, albeit with a declining regenerative capacity over time^19,20^. Acute total body irradiation induces rapid thymic involution^21–23^, primarily attributed to the severe and rapid reduction of thymocytes, particularly CD4 and CD8 double-positive immature thymocytes. Additionally, a body of literature indicates that TECs are also susceptible to the effects of irradiation and undergo homeostatic recovery^23–27^.

Previous studies provided cellular and molecular mechanistic insights on the recovery of TECs from aging-independent involution of the thymus. Several cell types appear to be involved in the TEC recovery. Type 3 innate lymphoid cells (ILCs) were engaged in TEC proliferation after irradiation^23^. Moreover, the contribution of eosinophils, natural killer T cells, and type 2 ILCs (ILC2) was found^18,26,27^. In addition to these hematopoietic origin cells, fibroblasts and thymic mesenchyme are reportedly necessary for regeneration during irradiation-induced thymic involution^18,27^. In the molecular mechanisms, some cytokines and growth factors appear to be involved in the thymic regeneration: IL-22^23^, IL-33^27^, type 2 cytokines^18,26^, receptor activator of nuclear factor κ B ligand (RANKL)^28^, endothelial-induced bone morphogenetic protein 4^29^, and fibroblast growth factor 7 (FGF7)^30^ reportedly contribute to the recovery from aging-independent thymic involution. In addition to these positive regulators, TGF-β signaling in TECs suppresses thymic recovery in the early phase after irradiation^31^.

Notably, previous studies reported a bimodal effect of acute total body irradiation on the murine thymus^18,32,33^. Initially, irradiation induced an acute and severe reduction in cell number within 3 to 4 days post-irradiation, followed by a subsequent recovery to levels comparable to those in control specimens. Following this, the thymic cell number decreased again, eventually leading to a gradual recovery. This bimodal reduction phenomenon was observed in thymocytes^32^. Nevertheless, the precise effect of this “second impact” on the frequency and properties of TECs remains to be elucidated.

In this study, we analyzed TECs of mice exposed to acute sublethal total body irradiation at three timepoints: 5 days post-irradiation, corresponding to the initial impact of the irradiation, 15 days post-irradiation, corresponding to the recovery phase from the initial impact, and 30 days post-irradiation, representing the second impact. We found that the second impact caused a preferential reduction in mTECs expressing high levels of CD80. Furthermore, RNA sequencing analysis showed increased expression of gene sets associated with tuft-like TECs and reduced expression of a gene signature of Aire-expressing mTECs in mature mTECs. Consequently, the second impact of acute irradiation occurs not only on thymocytes but also on TEC frequency and properties.

## Result

### mTECs expressing high levels of CD80 were preferentially reduced due to the second impact of acute irradiation

To assess the long-term effects of acute (a high dose delivered in a short period) sublethal total body irradiation on thymic cell numbers, we analyzed thymic tissues from 7-week-old C57BL/6 mice at 5, 15, and 30 days post-irradiation (5.5 Gy). Consistent with previous reports^23–25,28,32^, the total thymic cell counts were reduced at Day 5 post-irradiation compared to age-matched unirradiated controls (Figure 1A). On Day 15, total thymic cell counts did not significantly differ between irradiated and unirradiated thymuses, indicating recovery of thymic cells following the initial impact of acute irradiation (Figure 1A). However, at 30 days post-irradiation, total thymic cell counts were significantly reduced again compared to unirradiated controls (Figure 1A). Hence, a second impact of acute irradiation on the thymus occurred between 15 and 30 days post-irradiation in this experimental setting.

**Figure 1.**
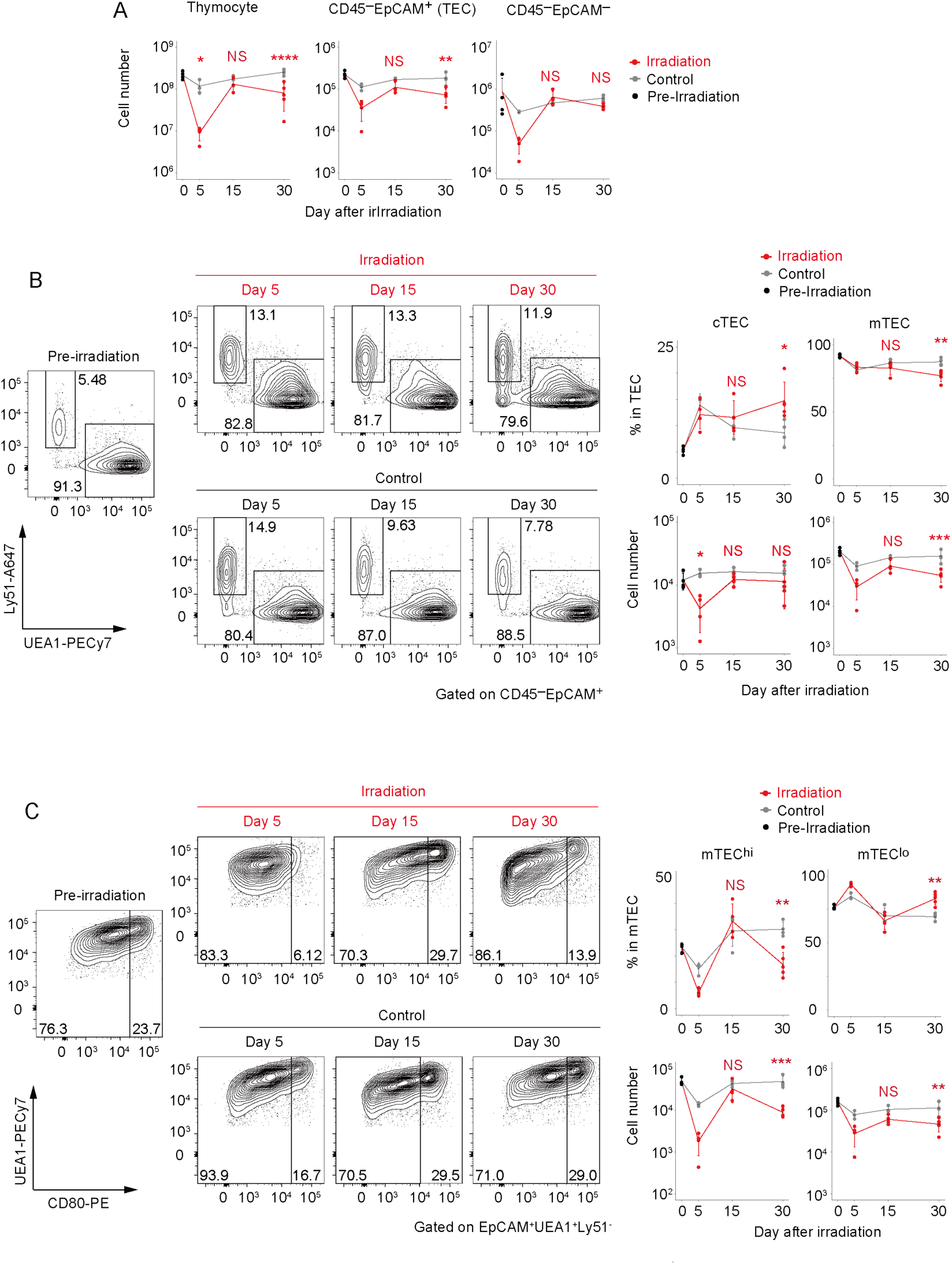
Long-term influence of acute sublethal irradiation on thymic cells. A. Number of thymocytes, CD45^−^EpCAM^+^ cells, and CD45^−^EpCAM^−^ cells after sublethal irradiation. Red indicates samples and gray indicates age-matched control. N = 4 for pre-irradiation, Day 5, Day 15, and control Day 30. N = 5 for radiation Day 30. One-way ANOVA with Tukey’s multiple comparisons test. * P < 0.05, ** P < 0.01, and **** P < 0.0001. NS indicates not significant. B. Flow cytometric analysis of TECs. Ly51^+^UEA-1^−^TECs and Ly51^−^UEA-1^+^TECs are designated as cTECs and mTECs, respectively. The changes in cell numbers and percentages of cTECs and mTECs within the total thymic epithelial cells (TECs; CD45^−^EpCAM^+^) are summarized in the graphs. N = 4 for pre-irradiation, Day 5, Day 15, and control Day 30. N = 5 for radiation Day 30. Two-way ANOVA test. * P < 0.05, ** P < 0.01, and *** P < 0.001. NS indicates not significant. C. Flow cytometric analysis of mTECs. mTECs were separated based on their expression levels of CD80 into mTEC^hi^ (high CD80 expression) and mTEC^lo^ (low CD80 expression). The changes in cell numbers and percentages of mTEC^hi^ and mTEC^lo^ in the total mTECs are summarized in the graphs. N = 4 for pre-irradiation, Day 5, Day 15, and control Day 30. N = 5 for radiation Day 30.One-way ANOVA with Tukey’s multiple comparisons test. ** P < 0.01 and *** P < 0.001. NS indicates not significant.

We investigated the second impact of acute irradiation on thymic epithelial cells (TECs) using flow cytometric analysis. Comparative analysis with age-matched controls revealed a significant reduction in the cell number of TECs (EpCAM^+^CD45^−^) on Day 30, but not on Day 15 (Figure 1A and Supplementary Figure 1). In contrast, the cell number of EpCAM^−^CD45^−^ stromal cells, predominantly composed of fibroblasts and endothelial cells, remained stable throughout this period. Consequently, our findings indicate the susceptibility of TECs, alongside lymphocytes as reported ^32^, to a second impact of acute radiation exposure.

TECs were further separated into medullary TECs (mTECs) and cortical TECs (cTECs) based on the expression of surface markers Ly51 (cTEC marker) and the binding of Ulex europaeus lectin 1 (UEA-1; mTEC marker). Despite the reduction in the total number of TECs by Day 5 post-irradiation, the ratios of mTECs (UEA-1^+^Ly51^−^) and cTECs (UEA-1^−^Ly51^+^) in the total TEC population remained consistent with those observed in age-matched controls (Figure 1B). On Day 15, there were no significant differences in the cell number and percentage of mTECs and cTECs, consistent with the recovery from the initial impact observed on Day 15. Following this recovery, the ratio of mTECs in the total TEC population decreased, while that of cTECs increased on Day 30 (Figure 1B). In terms of cell numbers, mTECs showed a significant reduction, whereas cTECs remained relatively stable (Figure 1B). These data suggest that the secondary impact of acute irradiation preferentially affects mTECs, leading to a greater reduction in mTEC cellularity compared to cTECs.

mTECs are categorized into two major subpopulations based on the expression levels of the co-stimulatory molecule CD80 and MHC class II. mTECs expressing high levels of CD80 (mTEC^hi^) are considered a mature type of mTEC, while mTECs expressing low levels of CD80 (mTEC^lo^) mainly encompass immature mTECs. In the initial impact on Day 5, the numbers of both mTEC^hi^ and mTEC^lo^ were reduced, with the reduction being more pronounced in mTEC^hi^ than in mTEC^lo^ (Figure 1C). On Day 15 post-irradiation, the reduction in the numbers of these mTEC subsets did not significantly differ between irradiated and unirradiated mice (Figure 1C), suggesting their recovery. Notably, on Day 30, the numbers of both mTEC^hi^ and mTEC^lo^ were reduced again (Figure 1C). The ratio of mTEC^hi^ in total mTECs was reduced, suggesting a significantly more pronounced impact on the mTEC^hi^ subset, while both mTEC^hi^ and mTEC^lo^ populations were affected by a second acute radiation exposure.

### The frequency of subpopulations in mature mTEC was not largely affected by the second impact of the irradiation

mTEC^hi^ consists of some subpopulations exhibiting unique gene expression profiles. To further characterize the second impact of acute irradiation on the mTEC^hi^ subpopulation, we examined the expression of surface markers Sca-1 and CD24 in this population ^34^. A previous study has delineated the differentiation process of mature mTECs in the order of CD24^−^Sca-1^−^, CD24^+^Sca-1^−^, and CD24^+^Sca-1^+^ mTEC^hi^, accompanied by changes in gene expression, notably decreased Aire expression and increased genes expressed in some post-Aire mimetic TECs^34^. Flow cytometric analysis revealed a significant reduction in the CD24^−^Sca-1^−^ cell fraction compared to other fractions in the mTEC^hi^ subpopulation on Day 5 (Figure 2), with recovery observed on Day 15. On Day 30, the proportion of cells in the CD24^−^Sca-1^−^ fraction had slightly decreased, while the proportion in the CD24^+^Sca-1^−^ fraction had increased. The proportion of cells in the CD24^+^Sca-1^+^ fraction remained constant. Despite these proportion changes, the absolute cell numbers were similarly reduced across all subsets (Figure 2). These findings suggest that, following an initial severe reduction in the relatively early stage of mTEC^hi^ and its subsequent recovery, the secondary impact of acute irradiation minimally affects the post-differentiation process of mature mTECs and might predominantly influence the differentiation step from immature mTECs to mature mTECs.

**Figure 2.**
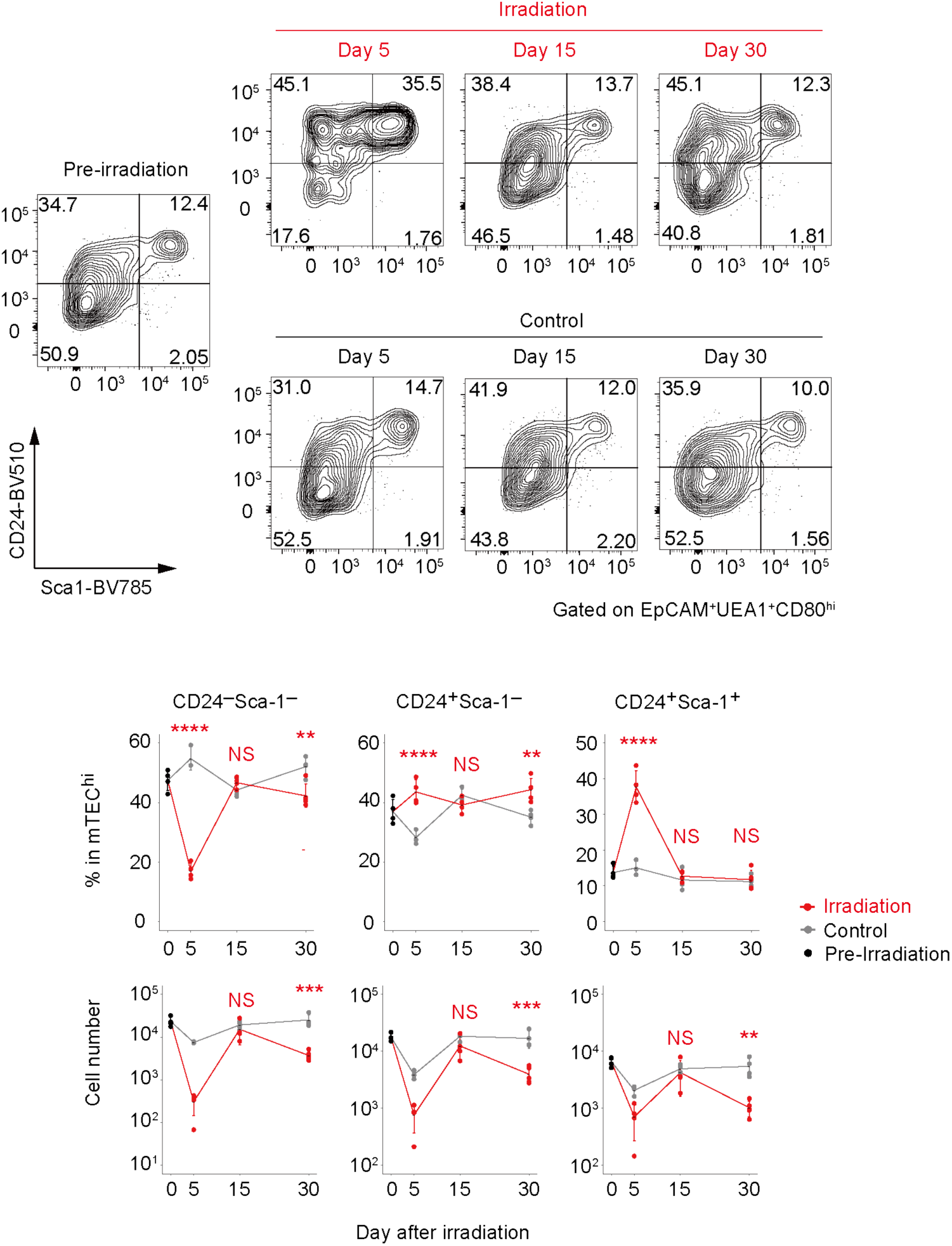
Influence of acute sublethal irradiation on mature mTECs. Flow cytometric analysis of mTEC^hi^ subpopulations. mTEC^hi^ was separated by expression levels of CD24 and Sca-1. The changes in cell numbers and percentages of CD24^−^Sca-1^−^, CD24^+^Sca-1^−^ and CD24^+^Sca-1^+^ fractions in the total mTEC^hi^ are summarized in the graphs. N = 4 for pre-irradiation, Day 5, Day 15, and control Day 30. N = 5 for radiation Day 30.One-way ANOVA with Tukey’s multiple comparisons test. ** P < 0.01, *** P < 0.001 and **** P < 0.0001. NS indicates not significant.

### The second impact of the irradiation leads to a change of gene expression profile in some mature mTEC subsets

We next aimed to investigate the second impact of acute irradiation on the gene expression. Considering the influence of the second impact on the differentiation step from immature to mature mTECs, we investigated the gene expression of CD24^−^Sca-1^−^ and CD24^+^Sca-1^−^ mTEC^hi^, which are in the relatively early stages of differentiation in mature mTECs. These mature mTEC subsets were sorted at pre-irradiation (Day 0), and at 15 and 30 days post-irradiation, along with respective age-matched control samples, for RNA-seq analysis. Data analysis indicated that the expression level of Aire was significantly reduced, while the expression of marker genes associated with post-Aire mTECs was elevated in the CD24^+^Sca-1^−^ mTEC^hi^ subset compared to the CD24^−^Sca-1^−^ mTEC^hi^ subset (Supplementary Figure 2). These observations were consistent with previously reported gene expression changes associated with mTEC differentiation^34^.

Moreover, principle component analysis (PCA) of the RNA-seq data revealed a clear separation between CD24^−^Sca-1^−^ and CD24^+^Sca-1^−^ mTEC^hi^ along the PC1 axis (Figure 3A), indicating distinct gene expression profiles between these subsets. Notably, both CD24^−^Sca-1^−^ and CD24^+^Sca-1^−^ mTEC^hi^ on Day 30 are separated from the other mTECs in each group along the PC2 axis (Figure 3A), implying the second impact of acute irradiation in the gene expression profile of these mTEC subsets on Day 30.

**Figure 3.**
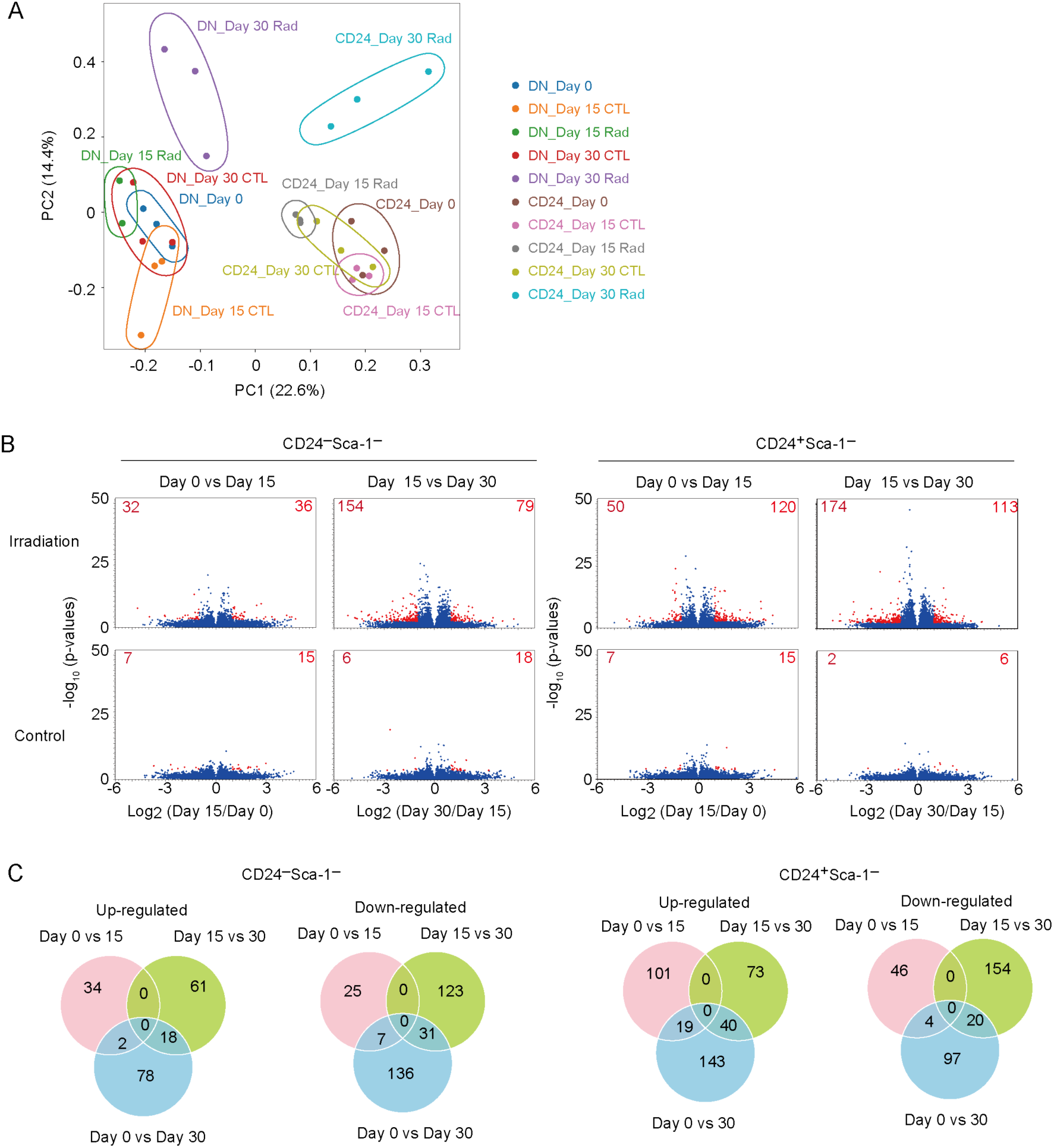
Gene expression change in mature mTEC subpopulations by the second impact of acute sublethal irradiation. A. PCA analysis of RNA-seq data. “DN” indicates CD24^−^Sca-1^−^mTEC^hi^, “CD24” indicates CD24^+^Sca-1^−^mTEC^hi^. “Rad” indicates irradiation samples and “CTL” indicates age-matched control. N = 3 for all samples except for N = 2 for DN_Day 15 Rad. B. Volcano plots for the RNA-seq data of CD24^−^Sca-1^−^mTEC^hi^ and CD24^+^Sca-1^−^mTEC^hi^ populations after radiation, compared to age-matched controls. Differentially expressed genes between Day 0 and Day 15 (left) and between Day 15 and Day 30 (right) are displayed. Red indicates differentially expressed genes (2-fold change and FDR P < 0.05). N = 3 for all samples except for N = 2 for CD24^−^Sca-1^−^ mTEC^hi^ irradiation sample. C. Venn diagram of differentially expressed gene sets.

Further data analysis revealed significant alterations in gene expression between pre-irradiation (Day 0) and Day 15 post-irradiation in both CD24^−^Sca-1^−^ and CD24^+^Sca-1^−^ mTEC^hi^ populations, suggesting a sustained effect of acute irradiation at the transcriptional level (Figure 3B), despite minimal changes in cell numbers as observed in flow cytometric analysis. From Day 15 to Day 30, additional shifts in gene expression profiles were observed in both CD24^−^ Sca-1^−^ and CD24^+^Sca-1^−^ mTEC^hi^ subsets (Figure 3B). Notably, differentially expressed gene sets between Day 15 and Day 30 entirely differ from those between pre-irradiation (Day 0) and Day 15 (Figure 3C and Supplementary File 1). This indicates that the gene expression changes from Day 15 to Day 30 were not simply due to the sustained effect of the initial impact but ascribed to the second impact. Consistently gene sets up-regulated from Day 15 to Day 30 remained relatively stable between Day 0 and Day 15 (Figure 4). Conversely, gene sets down-regulated during this period tended to be up-regulated between Day 0 and Day 15 (Figure 4). This implies that the second impact may counteract the sustained effect of the initial impact on the gene expression profile.

**Figure 4.**
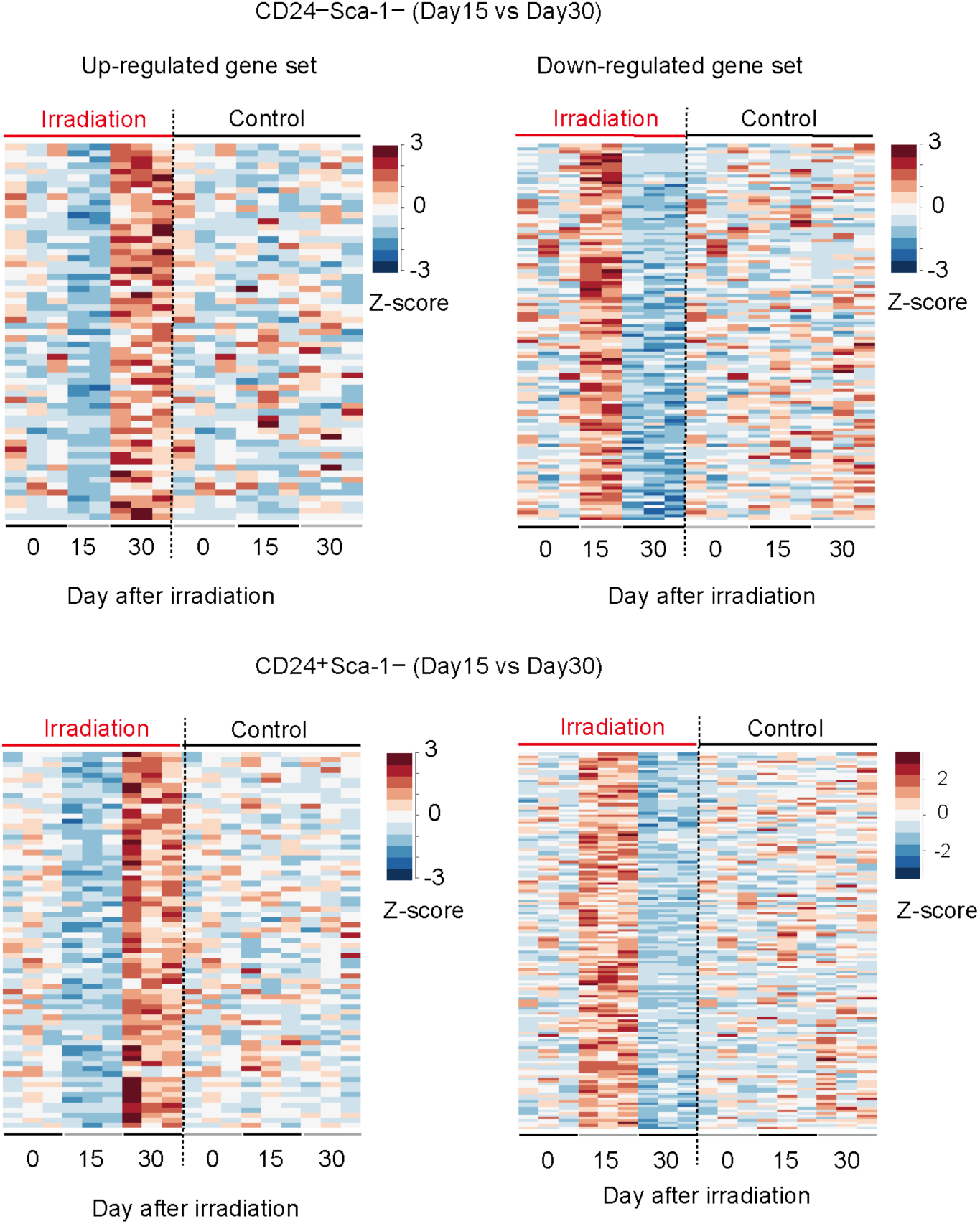
Heat maps of z-scores for the expression level of up-regulated and down-regulated gene sets from Day 15 to Day 30 in CD24^−^Sca-1^−^ mTEC^hi^ (upper panels) and CD24^+^Sca-1^−^ mTEC^hi^ (lower panels).

Given that CD24^−^Sca-1^−^ and CD24^+^Sca-1^−^ mTEC^hi^ contain Aire^+^ mTECs and Post-Aire mimetic mTECs, we analyzed the expression of signature genes for these cell types. Since Post-Aire mimetic mTECs are classified into several subsets, we first obtained marker gene sets for these mTEC subpopulations through re-analysis of a previous dataset ^35^. Specifically, after clustering and UMAP analysis of the dataset, each cluster was assigned according to the cell classification defined by the previous study ^35^. Following this assignment, genes exhibiting more than a 4-fold change and statistically significant (FDR p-value < 0.05) were designated as marker genes for each respective mTEC subset (Supplementary Figure 3 and Supplementary File 2).

For each marker gene set, gene expression changes from Day 15 to Day 30 in CD24^−^Sca-1^−^ and CD24^+^Sca-1^−^ mTEC^hi^ were plotted (Figure 5 and Supplementary Figure 4). The data showed that, from 15 to 30 days after the irradiation, the down-regulated genes in both CD24^−^Sca-1^−^ and CD24^+^Sca-1^−^ mTEC^hi^ included a significantly higher number of genes expressed in AIRE-positive cells. Moreover, the up-regulated genes in CD24^+^Sca-1^−^ mTEC^hi^ included a higher number of genes expressed in tuft-like mTECs (Figure 5). For other mTEC subset gene signatures, while some individual genes in the marker gene sets exhibited significant variation, the overall changes in gene expression across the entire marker gene sets were minimal (Supplementary Figure 4).

**Figure 5.**
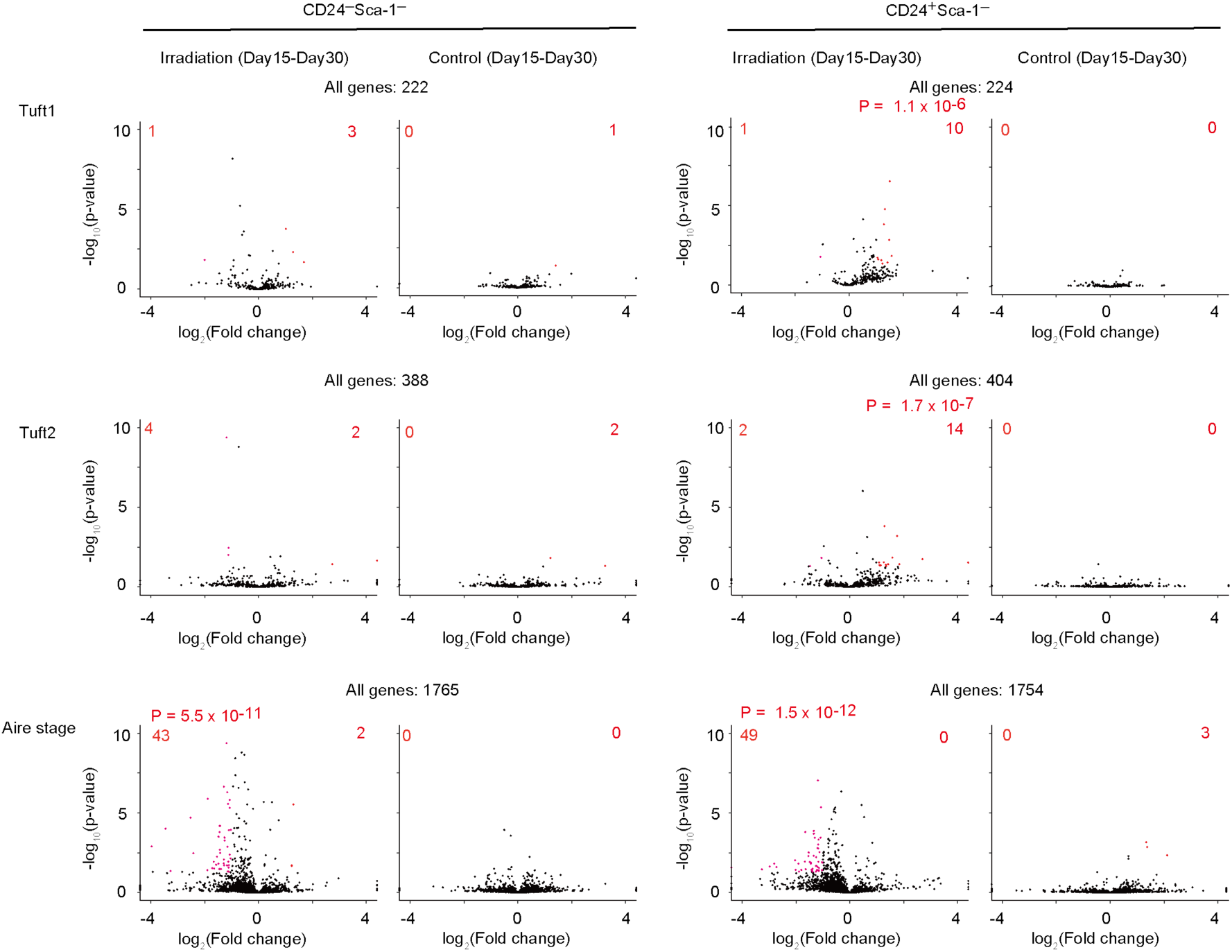
Volcano plots for expression data of marker genes for two tuft-like mTEC subsets and Aire mTEC in CD24^−^Sca-1^−^mTEC^hi^ and CD24^+^Sca-1^−^mTEC^hi^. Genes with increasing or decreasing expression levels were analyzed using a hypergeometric distribution to assess whether the corresponding gene sets were significantly enriched, with significance determined by the p-value.

### The second impact does not influence the cell number of CD80^−^Ly51^lo^UEA-1^+^TEC

Data suggested the influences of the second impact on the differentiation step from immature mTECs and mature mTECs. Since mTEC^lo^ encompasses immature mTECs, we analyzed the mTEC^lo^ population in detail. In addition to the reduction in cell numbers of mTEC^lo^ subsets during the second impact (Figure 1C), the surface expression level of CD80 protein appeared to be persistently decreased from the initial impact phase (Figure 6A). Moreover, the ratio of Ly51^lo^ cells increased in the mTEC^lo^ fraction post-irradiation (Figure 6B). In the cell number, whereas the Ly51^−^ mTEC^lo^ subsets showed a slight decrease on Day 30 compared to the age-matched control, the Ly51^lo^CD80^lo^UEA-1^+^ TECs remained largely unchanged during this period (Figure 6B), suggesting potential resistance to acute irradiation.

**Figure 6.**
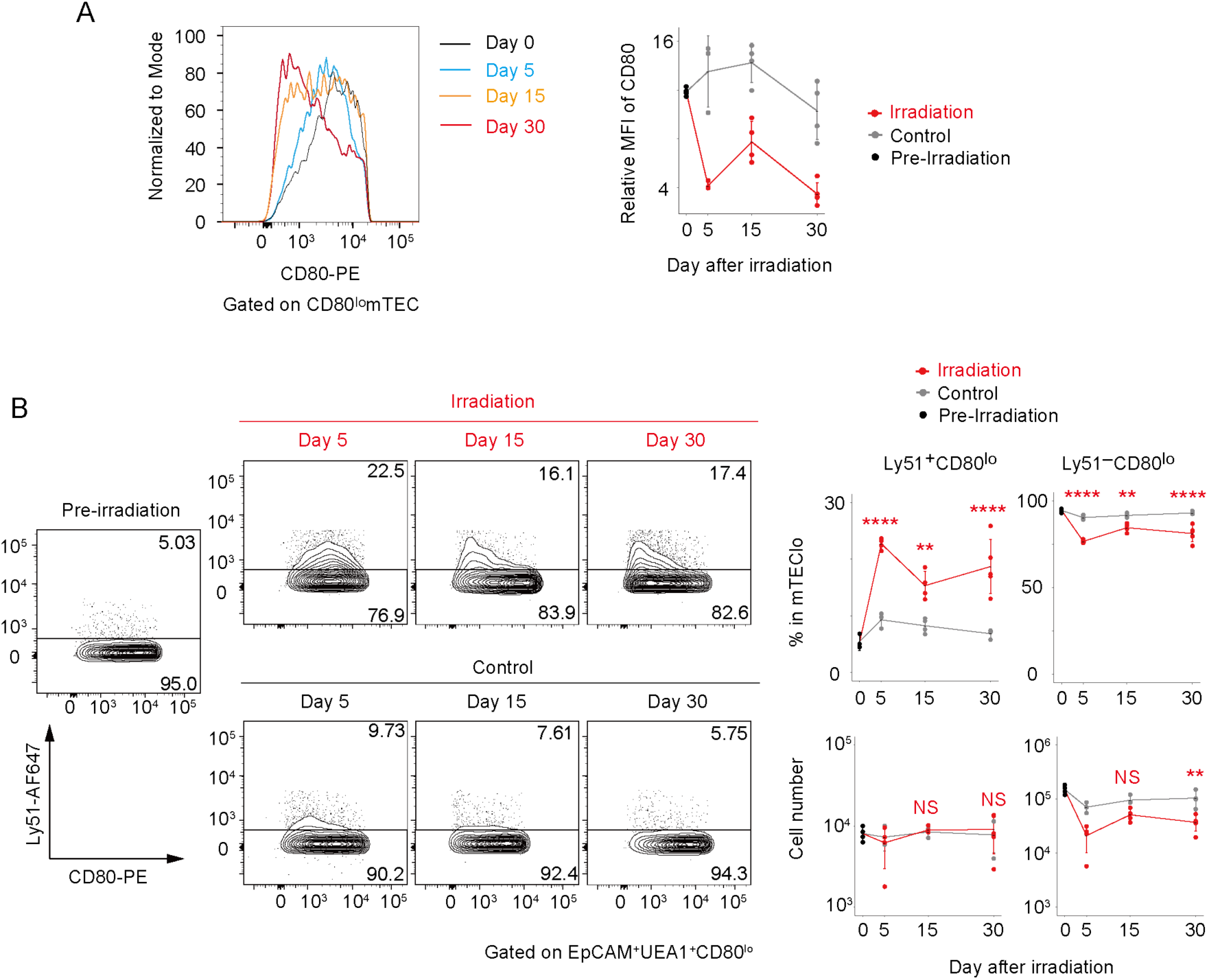
Expression of CD80 and Ly51 in mTEC^lo^ subset after acute irradiation. A. Flow cytometric analysis for assessing the expression level of CD80 in the mTEC^lo^ subset following irradiation. The relative median fluorescence intensities (MFIs) of CD80 in mTEC^lo^, compared to those in cTECs (used as the internal control), are plotted in the figure on the right. B. Flow cytometric analysis for Ly51^lo^ in mTEC^lo^ subsets. The changes in cell numbers and percentages of Ly51^lo^ cells and Ly51-negative cells in the mTEC^lo^ subset are summarized in the graphs.

## Discussion

Similar to a bimodal effect of acute sublethal irradiation on thymocyte reduction in mice^33^, our investigation indicates a similar bimodal effect on TECs, particularly mature mTECs. Given that mature mTEC development is reportedly promoted by cells of hematopoietic origin such as CD4 single-positive thymocytes via TNF family cytokine signaling^36,37^. The reduction of thymocytes may indirectly impair the homeostatic maintenance of mature mTECs during the second impact. However, because the initial impact is sustained in the mTEC^lo^ and gene expression profile in mature mTECs, it is possible that the combination of the indirect effect by lymphocytes and TEC-intrinsic mechanism results in the second effect on TECs. Moreover, it should be noted that the cell number of mTEC^lo^ was further reduced from Day 15 to Day 30. Thus, another mechanism controlling the differentiation of immature mTECs from their progenitor may be impaired during the second impact. This is also an important issue to be addressed in the future.

We noted that the cell number of UEA-1^+^Ly51^lo^CD80^−^ fraction was maintained on Day 15 and Day 30 after acute irradiation. Although Ly51 is considered to be a marker for cortical TECs, a previous study showed that Ly51^+^UEA-1^+^ TECs are progenitors of mTECs in the embryonic thymus^38^. Thus, a similar type of mTEC progenitors may be radio-resistance and supply their progeny of mTECs.

Previous studies revealed tuft-like mTECs were resistant to dexamethasone and the irradiation-inducing damage^18^. Together with fibroblasts, tuft-like mTECs activate ILC2 through IL-25 signaling. Our data suggested that tuft marker genes were up-regulated in the CD24^+^Sca-1^−^ mTEC subset on Day 30 as compared to Day 15. Thus, our data suggested a possibility that, similar to dexamethasone-induced thymic involution and the first impact of irradiation, tuft-like cells are resistant also to the second impact, thereby promoting the recovery of the thymus in the following recovery stage.

A previous study showed that bone marrow transplantation after total body irradiation provokes autoimmunity^39^. Our data suggested that the gene signature of Aire^+^ mTECs was reduced during the second impact. This suggests that thymic self-tolerance mechanism may be impaired in a bimodal manner. Thus, our data may imply that the second impact of total body irradiation may provoke increase in the risk of autoimmunity for a long period.

## Methods

### Mice

Animal experiments were performed in accordance with the Guidelines of the Institutional Animal Care and Use Committee of RIKEN, Yokohama Branch (2018-075). C57BL/6JJcl mice (CLEA Japan) were maintained in standard controlled conditions with a 12-h lighting cycle and access to chow and water ad libitum. Seven-week-old female mice received 5.5 Gy of γ-irradiation. Polymyxin B sulfate salt (10 KU/ml) (SIGMA P1004-1MU) and neomycin trisulfate salt hydrate (50 mg/ml) (SIGMA N1876-25G) was supplied with sterile drinking water after the irradiation.

### RNA-seq

mTEC subpopulations were sorted and lysed in 20µl Lysis Buffer (0.02% 2-ME in 2x TCL (Qiagen)) using BD FACSAria TM III cell sorter (BD). After purification of nucleic acids with SPRIselect (Beckman Coulter) and subsequent treatment with RNase inhibitor consisting of DW (Nacalai) and Rnasin plus (40U/µl) (Promega) and DNase I solution consisting of DW (Nacalai), 5x PrimeScript Buffer (TAKARA) and 1 U/µl DNase I (Thermo Fisher), first-strand cDNA synthesis master mix containing 5x PrimeScript Buffer (TAKARA), PrimeScript RT Enzyme mix I (TAKARA), 1mg/ml T4 gene 32 protein (NEB) in DW (Nacalai), 4 µM oligo (dT) 18 (Thermo Fisher) in DW (Nacalai), and 100 µM 1st NSR primer (SIGMA) was added to the purified nucleic acids samples for reverse transcription and mixed by gentle tapping. Then the mixed samples were incubated in a T100 TM Thermal Cycler (BioRad) at 4℃ for 10sec, (ramp up 0.1℃/sec to 25℃) 25℃ for 10min, (ramp up 0.1℃/sec to 30℃) 30℃ for 10min, (ramp up 0.1℃/sec to 37℃) 37℃ for 30min, (ramp up 0.1℃/sec to 50℃) 50℃ for 5min, 94℃ for 5min. The samples were then held at 4℃ or on ice until the next step. The first-strand cDNA synthesis master mix samples were added to second-strand synthesis master mix containing DW (Nacalai), 10x NEB Buffer2 (NEB), 10 mM each dNTP (NEB), 100 µM 2nd NSR primer (SIGMA), and Klenow Fragment (3’-5’ exo-) (NEB) and mixed by gentle tapping. The mixed samples were then incubated in a T100 TM Thermal Cycler (BioRad) at 4℃ for 10sec, 16℃ for 60min, 70℃ for 10min. The samples were then held at 4℃ or on ice until the next step. After cDNA synthesis and subsequent purification by SPRIselect (Beckman Coulter), sequencing library DNA was prepared using the Tn5 tagmentation-based method. Paired-end sequencing was performed using a HiSeq X TM Ten (Illumina). Sequence reads were quantified for annotated genes using CLC Genomics Workbench (Version 23.0.4; Qiagen).

### Flow cytometry and cell sorting

Flow cytometric analysis and cell sorting were performed using BD FACSAria TM (BD).

Thymic stromal cells were prepared by mincing murine thymi with razor blades, pipetting thymus fragments up and down to remove lymphocytes and then digesting the fragments with RPMI 1640 medium containing Liberase (Roche, 0.05 U/ml) plus DNase I (Sigma-Aldrich) via incubation at 37℃ for 12 min three times. Before cell staining, anti-mouse CD16/32 (Biolegend, Cat; 101302) was used as Fc Blocking. Thymic stromal cells were stained with the first anti-mouse antibody mixture containing CD45-APC-Cy7 (Biolegend, Cat; 103116), TER119-APC-Cy7 (Biolegend, Cat; 116223), EpCAM-FITC (Biolegend, Cat; 118208), Ly51-Alexa Fluor 647 (Biolegend, Cat; 108312), UEA1-Biotine (Vector Laboratories, Ref; B-1065), CD80-PE (Biolegend, Cat; 104708), CD24-BV510 (Biolegend, Cat; 101831), and Sca1-BV785 (Biolegend, Cat; 108139) and second antibody mixture containing Streptavidin-PE-Cy7 (Biolegend, Cat; 405206) on ice for 20minutes each. Dead cells were excluded by 7-aminoactinomycin D staining. Data were analyzed using FlowJo TM 10 (Version 10.6.2).

### Statistical analysis

Statistical significance between mean values was determined using one-way ANOVA test.

## Acknowledgment

This work was supported by Grants-in-Aid for Scientific Research from JSPS (21K19391 and 23K27399 to T.A., 23K06385 to N.A., and 24K18386 to T.M) and by CREST from the Japan Science and Technology Agency (JPMJCR2011 to T.A.). The authors declare no competing financial interests.

## Author contributions

**Kano Namiki**, Data curation, Formal analysis, Investigation, Validation, Writing; **Wataru Muramatsu**, Data curation, Formal analysis, Investigation, Validation; **Takahisa Miyao**, Investigation, Validation; **Hiroto Ishii**, Formal analysis, Investigation; **Rin Endo**, Investigation, Validation**; Naho Hagiwara**, Data curation; **Maki Miyauchi**, Investigation, Validation; **Masaki Yoshida,** Investigation, Validation; **Nobuko Akiyama,** Supervision, Formal analysis, Investigation, Writing – review and editing;**Taishin Akiyama,** Funding acquisition, Investigation, Project administration, Supervision, Validation, Writing.

## Supplementary Figure and legends

**Supplementary Figure 1.**
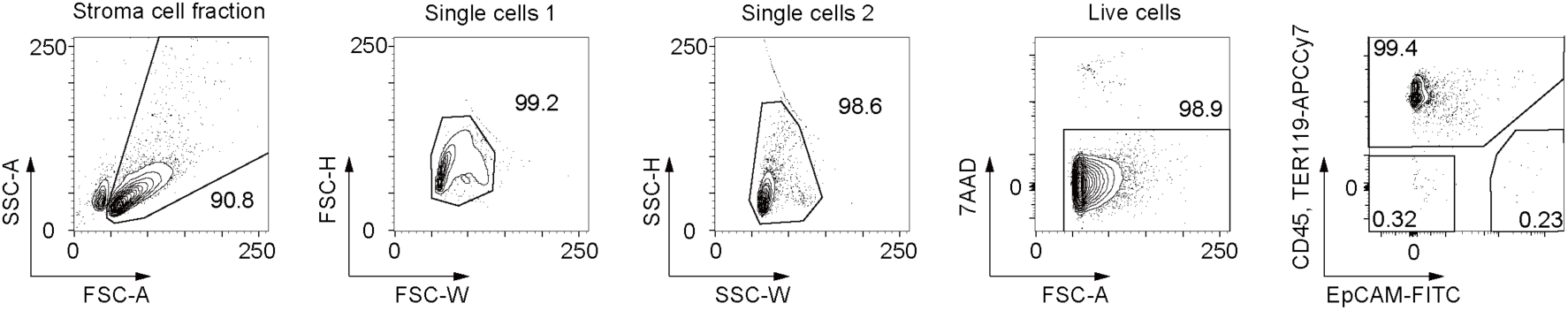
Gating strategy for TEC separation. TECs are defined as EpCAM^+^CD45^−^TER119^−^ fraction in alive single-cell gating.

**Supplementary Figure 2.**
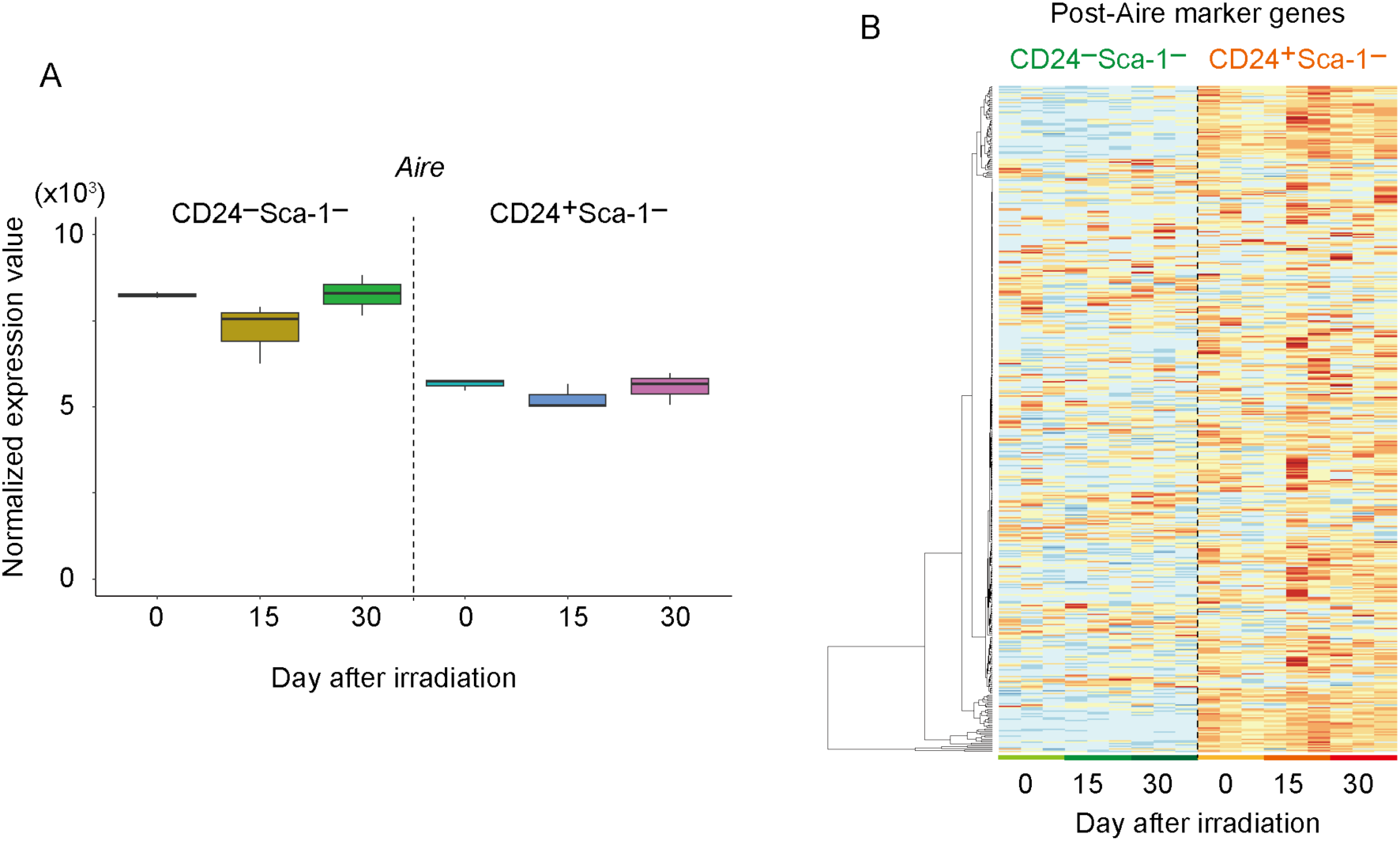
A. Normalized expression level of Aire in CD24^−^Sca-1^−^mTEC^hi^ and CD24^+^Sca-1^−^mTEC^hi^ B. Heatmap of post-Aire mimetic mTEC marker genes ^34^ in CD24^−^Sca-1^−^mTEC^hi^ and CD24^+^Sca-1^−^mTEC^hi^. Post-Aire marker genes used for analysis were reported previously by others (ref).

**Supplementary Figure 3.**
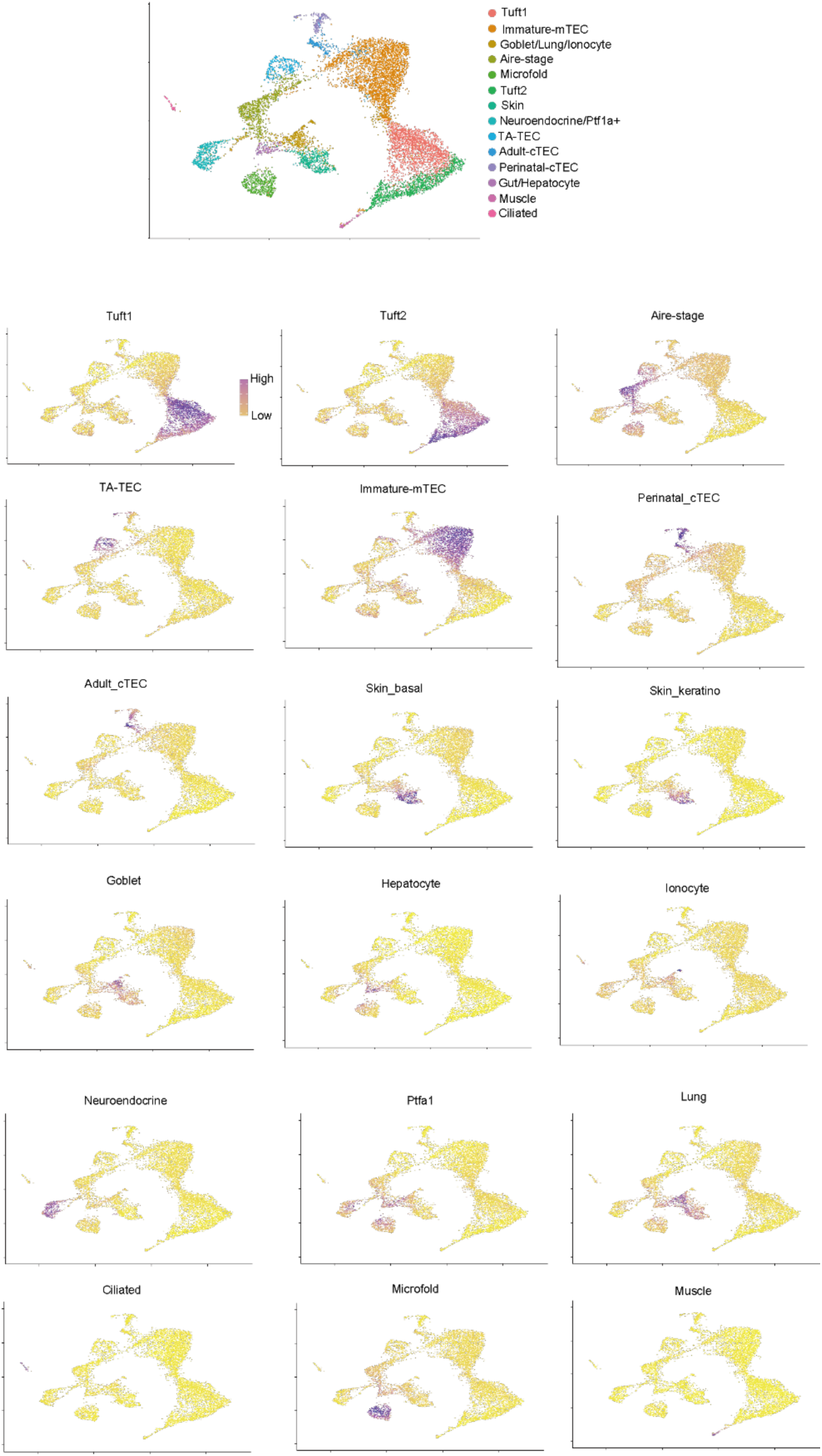
Re-analysis of scRNA-seq data of post-Aire mimetic cells. Each cluster was determined based on deposited data by using Seurat package, and cell types assigned in previous studies^35^ were shown for cluster assignment.

**Supplementary Figure 4.**
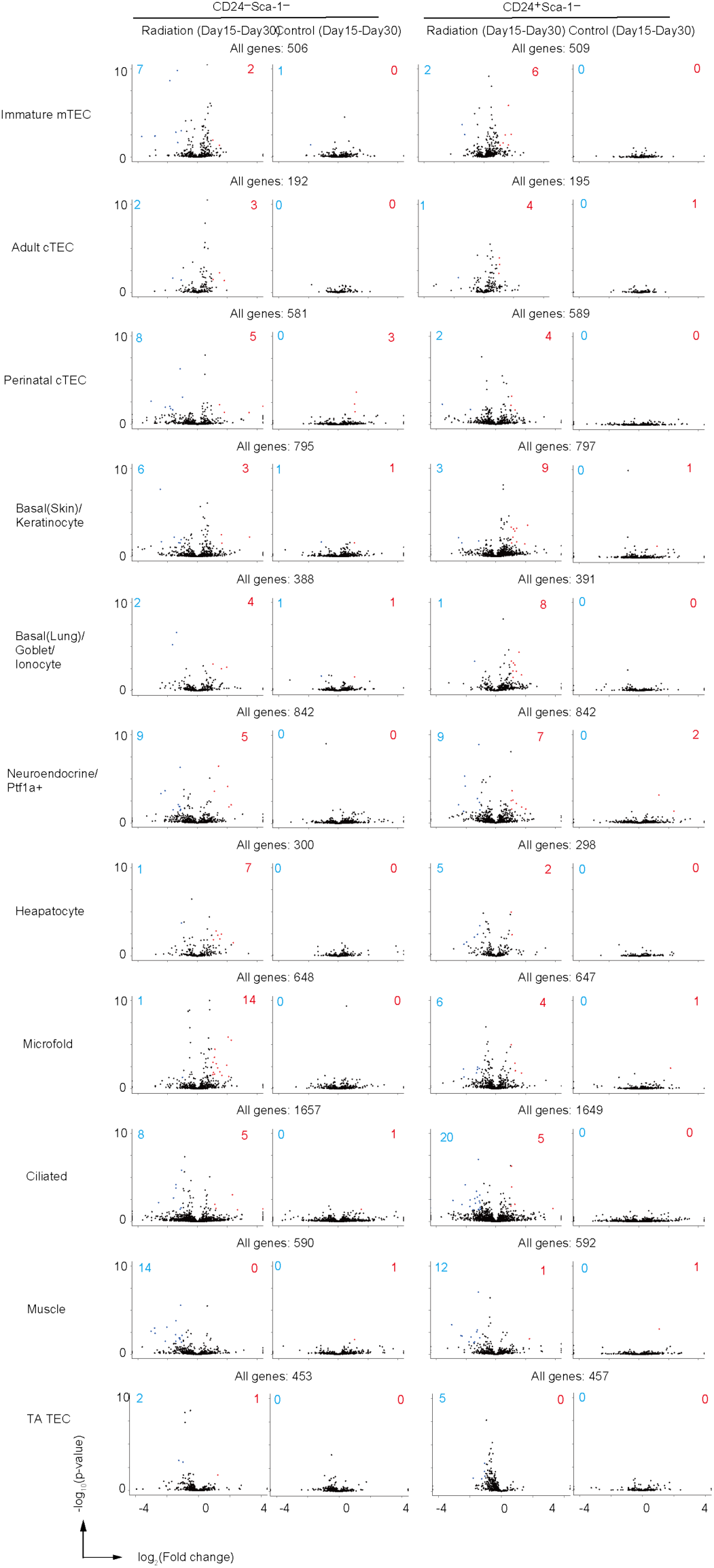
Volcano plots for expression data of marker genes for two tuft-like mTEC subsets, microfold mTECs, and Aire mTEC in CD24^−^Sca-1^−^mTEC^hi^ and CD24^+^Sca-1^−^mTEC^hi^

**Supplementary file 1**

Summary of differentially expressed gene sets.

**Supplementary file 2**

Lists of marker genes for mTEC subsets.

